# Task-related effective connectivity reveals that the cortical rich club gates cortex-wide communication

**DOI:** 10.1101/185603

**Authors:** Mario Senden, Niels Reuter, Martijn P. van den Heuvel, Rainer Goebel, Gustavo Deco, Matthieu Gilson

## Abstract

Higher cognition may require the globally coordinated integration of specialized brain regions into functional networks. A collection of structural cortical hubs - referred to as the rich club - has been hypothesized to support task-specific functional integration. In the present paper, we use a whole-cortex model to estimate directed interactions between 68 cortical regions from fMRI activity for four different tasks (reflecting different cognitive domains) and resting state. We analyze the state-dependent input and output effective connectivity of the structural rich club and relate these to whole-cortex dynamics and network reconfigurations. We find that the cortical rich club exhibits an increase in outgoing effective connectivity during task performance as compared to rest while incoming connectivity remains constant. Increased outgoing connectivity targets a sparse set of peripheral regions with specific regions strongly overlapping between tasks. At the same time, community detection analyses reveal massive reorganizations of interactions among peripheral regions, including those serving as target of increased rich club output. This suggests that while peripheral regions may play a role in several tasks, their concrete interplay might nonetheless be task-specific. Furthermore, we observe that whole-cortex dynamics are faster during task as compared to rest. The decoupling effects usually accompanying faster dynamics appear to be counteracted by the increased rich club outgoing effective connectivity. Together our findings speak to a gating mechanism of the rich club that supports fast-paced information exchange among relevant peripheral regions in a task-specific and goal-directed fashion, while constantly listening to the whole network.

## Introduction

The brain’s structural connectivity profile has been shown to contain a set of densely interconnected hub regions (Colizza, Flammini, Serrano, & Vespignani, 2006; van den Heuvel & Sporns, 2011; Zamora-López, Zhou, & Kurths, 2009). These regions, collectively termed the rich club, have been hypothesized to form a central high-capacity backbone for brain communication (van den Heuvel, Kahn, Goñi, & Sporns, 2012). As such they could play a crucial role in the integration of segregated brain regions into transient functional networks assumed to underlie higher cognition (Baars, 2005; Deco, Jirsa, & McIntosh, 2011; Deco, Van Hartevelt, Fernandes, Stevner, & Kringelbach, 2017; Dehaene & Naccache, 2001; Ghosh et al., 2008).

Several lines of research support this notion. Neuroimaging studies have shown that echoes of resting- and task related functional networks (Braga, Sharp, Leeson, Wise, & Leech, 2013; Leech, Braga, & Sharp, 2012) are present in the blood oxygen-level dependent (BOLD) time-series observed within cortical hub regions, indicative of the rich club serving as a central relay for cortical communication. Further support comes from the observation that selective disruption of connectivity among cortical hubs observed in schizophrenia is associated with reduced global communication capacity (Martijn P. van den Heuvel et al., 2013). Simulation studies have shown that cortical hubs, and specifically the rich club, may allow the brain to sustain a large functional repertoire characterized by diverse configurations of peripheral regions, i.e. regions of lower structural degree, around a stable high-degree core (Deco, Senden, & Jirsa, 2012; Senden, Deco, De Reus, Goebel, & van den Heuvel, 2014). Furthermore, there is evidence that rich club regions harmonize peripheral regions by engaging in infraslow oscillations which they exhibit exclusively during task performance (Senden, Reuter, van den Heuvel, Goebel, & Deco, 2017). Finally, the presence of hubs has been shown to increase network controllability (Liu, Slotine, & Barabási, 2011) by providing peripheral regions with the means to exhibit control over the network (Gu et al., 2015; Liu et al., 2011).

In light of these findings it is reasonable to conceive of the rich club as a central workspace of information integration wherein peripheral brain regions compete for control of the system, as recently proposed by Shanahan (2012). In this context, the rich club might act as a gate furthering communication among a winning set of task-relevant peripheral regions while intercepting signals from competing regions. During rest, the gate should largely be closed and the rich club should thus receive more input from peripheral regions than sending output to these regions (see figure 1A). During task performance, the gate would then be opened with the rich club increasing its output (figure 1B). This increase should not result in broad communication between all peripheral regions but rather in the establishment of a specific community with the rich club relaying information between a sparse, task-relevant, set of peripheral regions (figure 1C).

**Figure 1:**
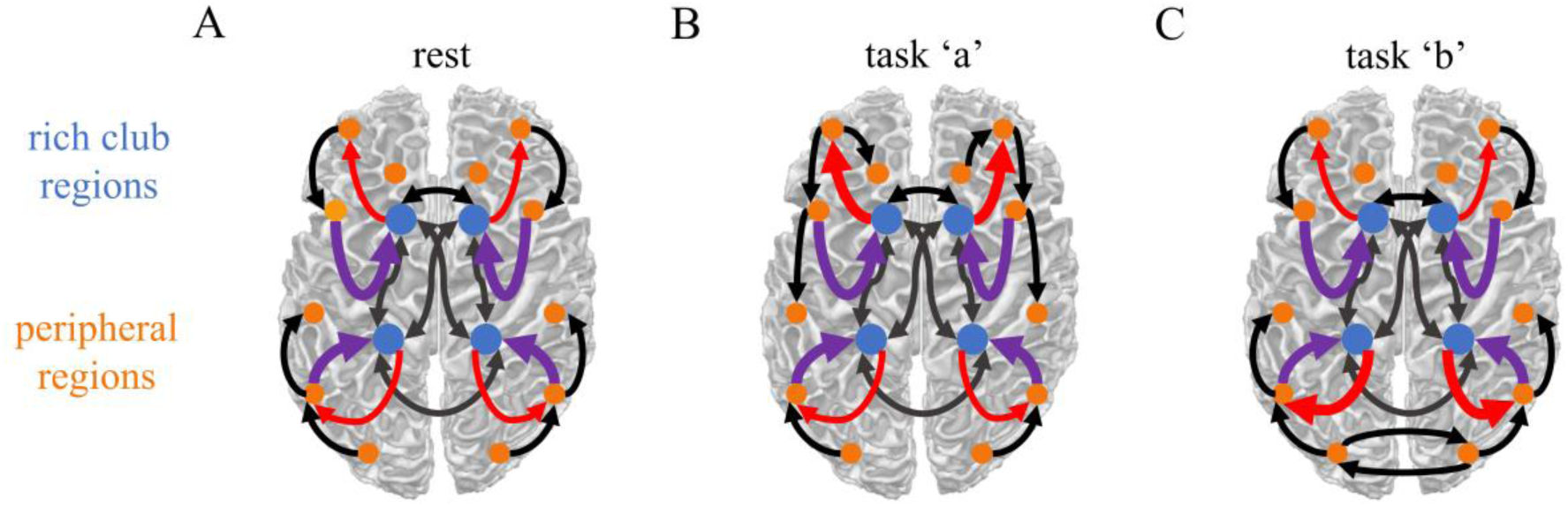
Brain regions are classified as rich club or peripheral regions. Communication occurs within each set of regions (black arrows) as well as between regions of different sets. *Panel A)* During rest, the rich club receives a fair amount of input from the periphery (purple arrows) but provides only little output back to peripheral regions (red arrows). *Panels B & C)* During task performance, the rich club increases the output it projects to the periphery. The specific peripheral targets of increased rich club output may depend on which task is currently being performed.

The present study aims to address these hypotheses using an integrative approach combining empirical functional magnetic resonance imaging (fMRI) data with large-scale computational modeling. Specifically, we employ a recently developed estimation procedure for whole-cortex effective connectivity (EC) in a noise-diffusion network model that reproduces the empirical spatiotemporal covariance of BOLD fluctuations (Gilson et al., 2016). The latter have been shown to convey information about the behavioral conditions of subjects (Mitra et al., 2015). Our whole-brain model captures changes in this spatiotemporal functional connectivity (FC) and interprets them in terms of local activity and network connectivity (Gilson et al., 2017). In contrast to previous modeling studies this procedure does not rely on identification of an optimal working point given fixed structural connectivity (Deco et al., 2013; Messé, Rudrauf, Benali, Marrelec, & Honey, 2014) but optimizes the strengths of interactions between brain regions in addition to their individual level of activity. A thorough comparison with previous models such as the dynamic causal model - or DCM (Friston, 2011) - and other techniques (e.g., partial correlations) is provided in Methods, to motivate our choice and highlight its advantages. Despite differences with DCM, we borrow the term “effective connectivity” from the DCM literature as our estimated connectivity describes the directional interactions between regions in the brain network. The EC matrix can be conceptualized as the transition matrix for the BOLD activities, for a cortical topology determined from the anatomical white-matter connections. The resulting estimates of directed EC allow us to study input/output relations of the rich club for rest and various task conditions in order to evaluate our gating hypothesis under an information propagation perspective. We optimized connectivity for a collection of fMRI measurements during which subjects engaged in a range of tasks tapping into different cognitive domains (visual working memory, response inhibition, mental rotation, verbal reasoning) and whose associated functional connectivity profiles are minimally overlapping (Smith et al., 2009).

## Materials and Methods

### Whole-Cortex Dynamic Model to Fit Empirical BOLD Covariances

The model comprised *N* = 68 cortical regions of interest ROI;(Desikan et al., 2006), whose local fluctuating activity (an amalgamation of internal processing and external input) is shaped by the recurrent effective connectivity matrix *C*. Following Gilson et al. (2016; https://github.com/MatthieuGilson/optimizationlinearECSigma.git), the model aims to reproduce BOLD covariances without and with time shift (see figure 2A), which are calculated as:

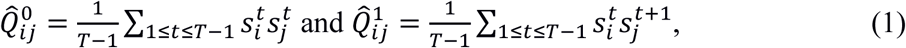

where 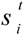 are the centered BOLD time series for each region 1 ≤ *i* ≤ *N* with time indexed by 1 ≤ *t* ≤ *T* (*T*=192 time points separated by a TR=2 seconds).

**Figure 2:**
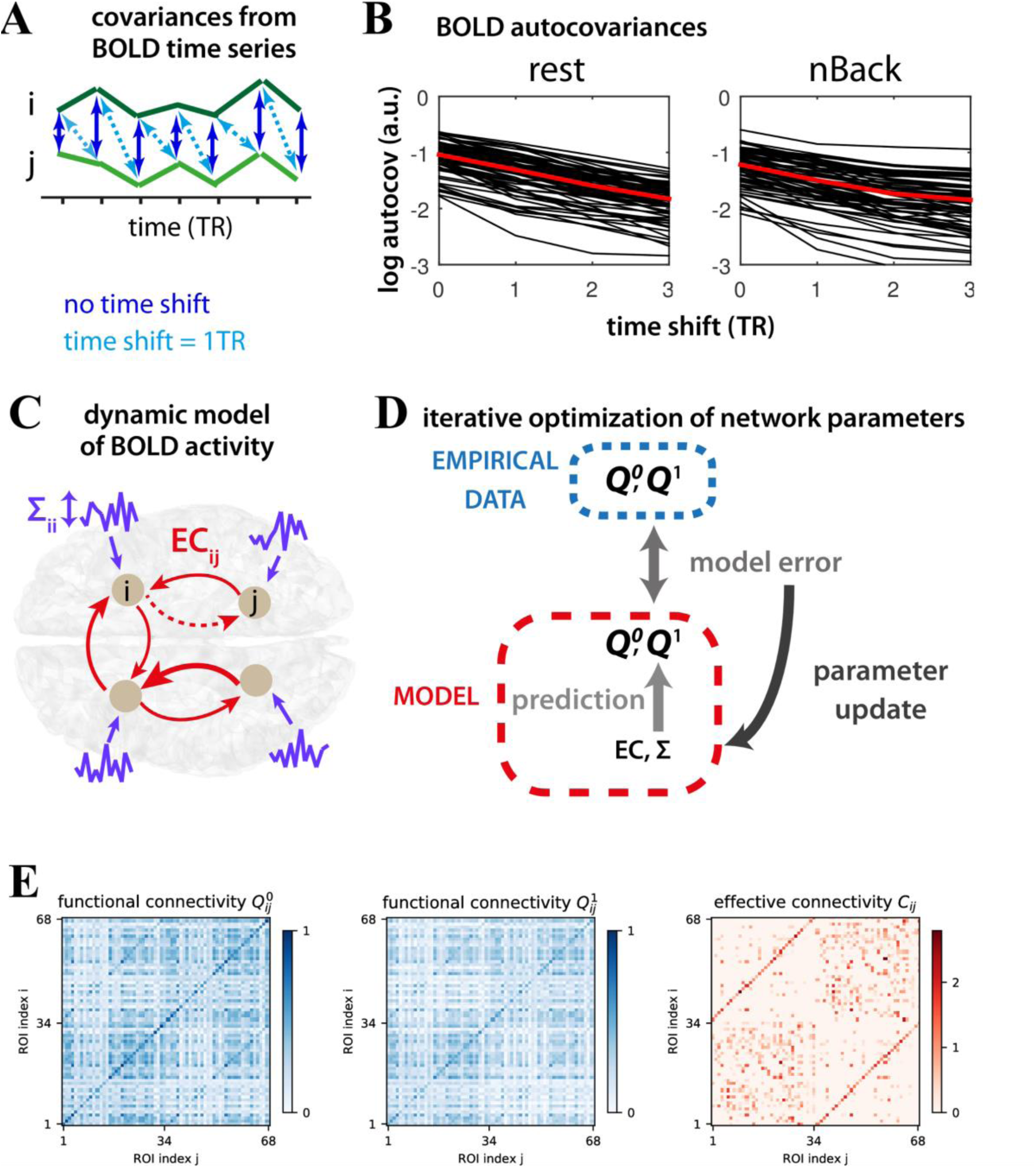
Schematic diagram of the model. *Panel A)* shows example BOLD time series (green) of two populations in the network. Dark blue arrows indicate temporal covariance with zero lag, whereas light blue arrows indicate covariance with time series *j* (light green) shifted by one TR (i.e. 2 seconds) with respect to time series *i* (dark green). *Panel B)* shows the logarithm of the autocovariances for each node as a function of time shift exemplary for rest and the n-back task. *Panel C)* shows a schematic diagram of the noise diffusion model: neural populations (gray circles) exhibit local, noisy, fluctuations (purple) as well as recurrent feedback (red arrows). Note that an existing connection (in the structural connectivity matrix) may end up with a zero weight (see dotted red arrow) which is equivalent to an absent connection. *Panel D)* shows a schematic representation of a single iteration within the Lyapunov optimization procedure. Model covariances (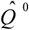 and 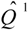) given current values of the network parameters EC and Σ are evaluated. From the comparison between model covariances with their empirical counterparts, the desired parameter updates for both parameters are calculated. *Panel E)* shows empirical covariance matrices *Q* ^0^ and *Q* ^1^ as well as effective connectivity matrix (C) resulting from the optimization procedure for resting state data. The cortical regions and their order can be found in table 1.

The local dynamics correspond to an Ornstein-Uhlenbeck process, where the nodal activity variable *x_i_* decays exponentially with time constant *τ_x_* and is affected by the rest of the network via 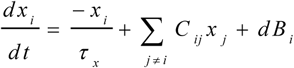. Here, each ROI experiences local fluctuations formally described by a Wiener process *dB_i_* (equivalent to white Gaussian noise) with a diagonal covariance matrix Σ. The between-region effective connectivity is embodied by the matrix *C*, whose skeleton is determined by a generic matrix of structural connectivity (see below). The advantage of this model is its analytical tractability, which allows for a quick calculation of its network covariance pattern *Q*^0^ and *Q*^1^. In essence, the model thus decomposes functional connectivity (FC) into two sets of parameters: effective connectivity and local variability. Those can be seen as a biomarker of brain dynamics, which captures the propagation of fluctuating BOLD activity across ROIs, as depicted schematically for 4 ROIs in figure 2C.

**Table 1:** Cortical regions.

| # | Region Name |
| --- | --- |
| 1 | left banks of the superior temporal sulcus |
| 2 | left caudalanterior-cingulate cortex |
| 3 | left caudalmiddle frontal gyrus |
| 4 | left cuneus |
| 5 | left entorhinal cortex |
| 6 | left fusiform gyrus |
| 7 | left inferior parietal cortex |
| 8 | left inferior temporal cortex |
| 9 | left isthmus–cingulate cortex |
| 10 | left lateral occipitalcortex |
| 11 | left lateral orbital frontal cortex |
| 12 | left lingual gyrus |
| 13 | left medial orbital frontal cortex |
| 14 | left middle temporal gyrus |
| 15 | left parahippocampal gyrus |
| 16 | left paracentral lobule |
| 17 | left pars opercularis |
| 18 | left pars orbitalis |
| 19 | left pars triangularis |
| 20 | left pericalcarine cortex |
| 21 | left postcentral gyrus |
| 22 | left posterior-cingulate cortex |
| 23 | left precentral gyrus |
| 24 | left precuneus |
| 25 | left rostralanterior cingulate cortex |
| 26 | left rostralmiddle frontal gyrus |
| 27 | left superior frontal cortex |
| 28 | left superior parietal cortex |
| 29 | left superior temporal gyrus |
| 30 | left supramarginal gyrus |
| 31 | left frontal pole |
| 32 | left temporal pole |
| 33 | left transverse temporal cortex |
| 34 | left insula |
| 35 | right banks of the superior temporal sulcus |
| 36 | right caudalanterior-cingulate cortex |
| 37 | right caudalmiddle frontal gyrus |
| 38 | right cuneus |
| 39 | right entorhinal cortex |
| 40 | right fusiform gyrus |
| 41 | right inferior parietal cortex |
| 42 | right inferior temporal cortex |
| 43 | right isthmus–cingulate cortex |
| 44 | right lateral occipitalcortex |
| 45 | right lateral orbital frontal cortex |
| 46 | right lingual gyrus |
| 47 | right medial orbital frontal cortex |
| 48 | right middle temporal gyrus |
| 49 | right parahippocampal gyrus |
| 50 | right paracentral lobule |
| 51 | right pars opercularis |
| 52 | right pars orbitalis |
| 53 | right pars triangularis |
| 54 | right pericalcarine cortex |
| 55 | right postcentral gyrus |
| 56 | right posterior-cingulate cortex |
| 57 | right precentral gyrus |
| 58 | right precuneus |
| 59 | right rostralanterior cingulate cortex |
| 60 | right rostralmiddle frontal gyrus |
| 61 | right superior frontal cortex |
| 62 | right superior parietal cortex |
| 63 | right superior temporal gyrus |
| 64 | right supramarginal gyrus |
| 65 | right frontal pole |
| 66 | right temporal pole |
| 67 | right transverse temporal cortex |
| 68 | right insula |

### Estimation Procedure for Effective Connectivity and Local Excitability

We iteratively tune model parameters *C* and *Σ* such that model covariance matrices *Q*^0^ and *Q*^1^ best reproduce the empirical FC 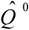 and 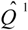. We summarize the essential steps of the procedure described in Gilson et al. (2016) that iteratively optimizes the network parameters *C* and *Σ* For each state (rest & tasks), we calibrate the model by calculating the time constant τ_x_ associated with the exponential decay of the autocovariance function 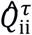 averaged over all regions: 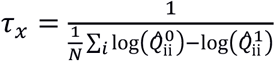 (corresponding to the average slope for the red curves in figure 2B). This gives the diagonal elements of the Jacobian of the dynamic system: 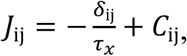 where *δ*_ij_ is the Kronecker delta. Starting from no recurrent connectivity (*C*=0), the model FC matrices *Q*^0^ and *Q*^1^ are calculated by solving the Lyapunov consistency equation JQ^0^+*Q*^0^*J*^*T*^+Σ=0 using the Bartels-Stewart algorithm and then evaluating *Q*^1^=*Q*^0^*e*^J^*T*^^ with the matrix exponential. The desired Jacobian update is the matrix *ΔJ^T^*=(*Q*^0^)^−1^[*ΔQ*^0^+*ΔQ*^1^*e*^−J^*T*^^], which involves the FC error between the empirical and model matrices: 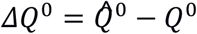 and 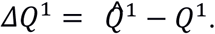 This yields the connectivity update *ΔC*_ij_=*η_c_ΔJ*_ij_ for existing connections and the local variance update ΔΣ_ii_=−*η_Σ_*(*JΔQ*^0^+*ΔQ*^0^*J^T^*)_ii_ for the diagonal elements only. We use *η_C_*=0.0001, *η_Σ_*=0.1, and impose non-negativity for the weights in *C* and the diagonal elements of *Σ*

Our version of EC relies on a linear-feedback model, which would correspond to the mean (linearized) effect in a network equipped with more elaborate nodal dynamics, allowing for the optimization of a large number of ROIs covering the whole cortex. In short, our EC lumps together local parameters modulating neuronal activity and synaptic interactions. The estimated changes in our EC thus describe the net effect resulting from possible compound causes. Therefore, the interpretation of the model is more phenomenological than with DCM. Nevertheless, to robustly estimate EC for many ROIs in a whole-brain context, the original DCM would require simplifications (Frässle et al., 2015), in line with our approach. Technically, our network model relies on a maximum-likelihood estimate of the whole-cortex EC, without the full Bayesian machinery used with DCM (Friston, 2011). The model parameters are tuned to reproduce the empirical cross-covariances between ROIs, which are canonically related to the cross spectral density used in recent studies that apply DCM to resting state fMRI data (Frässle et al., 2017; Friston, Kahan, Biswal, & Razi, 2014). Moreover, the proposed model relies on an exponential approximation of BOLD autocovariance (locally over a few TRs) and discards very slow-frequency fluctuations below 0.01 Hz. This model-based approach has been successfully applied to identify changes in the cortical coordination between rest and movie viewing (Gilson et al., 2017). In essence, our approach aims to bridge the gap between whole-brain modeling where the connectivity is fixed and taken from DTI (Deco et al., 2013; Messé et al., 2014) and studies focusing on a few cortical areas only to test hypotheses on specific brain subsystems (Goebel, Roebroeck, Kim, & Formisano, 2003; He, 2011).

In addition to the estimated EC, which can be seen as a matrix of partial correlations with information about directionality, the model estimation also involves input variances. This means that the changes in empirical functional connectivity are explained by two distinct sets of local and network parameters and hence distinguishes between the contribution of local processing (potentially in response to external inputs) and network communication. Moreover, the model uses structural connectivity (from DTI) to only estimate putative connections, discarding the rest; this reduces the number of parameters to estimate and improves the estimation robustness. Compared to phenomenological measures such as phase differences for all pairs of ROIs, the estimated EC weights have to “make sense” collectively, as the transition matrix between the ROI activities that generates model functional connectivity. In this way, observation noise will not affect the measures equally and independently across the ROI pairs but the estimate will correspond to a model with minimal global error with respect to the covariance matrices.

### Structural Connectivity (SC)

We obtained high-quality diffusion-weighted MRI data of 215 subjects from the Q3 release of the human connectome project HCP;(Glasser et al., 2013; Van Essen et al., 2012). Combined diffusion tensor and generalized q-sampling imaging (GQI) were used to fit complex fiber architecture, with white matter pathways reconstructed by means of streamline tractography (Romme, de Reus, Ophoff, Kahn, & van den Heuvel, 2017; Yeh, Wedeen, & Tseng, 2010)., The cortex was parcellated into 68 cortical regions according to the widely used Desikan-Killany (DK) atlas (Desikan et al., 2006; Klein & Tourville, 2012) forming the regions of the reconstructed anatomical connectome map. Details on these processing steps can be found in de Reus and van den Heuvel (2014). A weighted group-average structural connectivity matrix was generated by averaging streamlines over subjects and keeping only those entries which had positive values for at least 60% of subjects (de Reus & van den Heuvel, 2013). Finally, since Gaussian resampling (or taking the logarithm) of the data has previously been shown to enhance correspondence between diffusion tractography and in vivo animal tract-tracing measurements of anatomical connectivity (Martijn P. van den Heuvel et al., 2015), the data was resampled with a mean of 0.5 and a standard deviation of 0.15 (Honey et al., 2009).

### Functional MRI Data

We used resting and task-state BOLD time-series data from fourteen healthy subjects (8 females, mean age = 28.76) previously described by Senden *et al.* (2017). Briefly, data comes from five functional runs consisting of a resting-state measurement (eyes closed), four individual task measurements including a visual n-back (n=2) task (Kirchner, 1958), the Eriksen flanker task (Eriksen & Eriksen, 1974), a mental rotation task (Shepard & Metzler, 1971), and a verbal odd-man-out task (Flowers & Robertson, 1985). All runs comprise 192 data points with tasks being continuously performed during this period. For the n-back and flanker task, stimuli were presented at a rate of 0.5 Hz; for the mental rotation and odd-man out tasks they were presented at a rate of 0.25 Hz. Task sequence was counterbalanced across participants with the exception that the resting state functional run was always acquired first to prevent carry-over effects (Grigg & Grady, 2010). The data were acquired using a 3 Tesla Siemens Prisma Fit (upgraded Tim Trio) scanner and a 64-channel head coil. Initial preprocessing was performed using BrainVoyager QX (v2.6; Brain Innovation, Maastricht, the Netherlands). This includes slice scan time correction, 3D-motion correction, high-pass filtering with a frequency cutoff of .01 Hz, and registration of functional and anatomical images. Subsequently, using MATLAB (2013a, The MathWorks,Natick, MA), signals were cleaned by performing wavelet despiking (Patel & Bullmore, 2015) and regressing out a global noise signal given by the first principal component of signals observed within the cerebrospinal fluid of the ventricles. Next, voxels were uniquely assigned to one of the 68 cortical ROIs specified by the DK atlas and an average BOLD time-series was computed for each region as the mean time-series over all voxels of that region (preprocessed BOLD time series per ROI and state are publicly available at doi:10.5061/dryad.mc7pd). Finally, empirical lag FC matrices for each participant were given by the covariances of the z-normalized BOLD time series according to equation (1). Model fitting and all analyses were performed using group average lag FC matrices.

### Louvain Community Detection Method

We identified ROI communities within the EC that maximize the modularity of possible partitions of the network (Newman, 2006), using the implementation developed by Bondel et al. (2008). Modularity measures the excess of connections between ROIs compared to the expected values estimated from the sum of incoming and outgoing weights for the nodes (targets and sources, respectively). In practice, the method stochastically determines local communities and then iteratively aggregates ROIs to maximize the modularity at each step, until reaching a minimum. In addition, a resolution parameter (equal to 1 for the RBContribution method in https://pypi.python.org/pypi/louvain/) is used to enforce community coherence. The process was repeated 30 times for each task to average over the initial random clustering. In the end, the reported value is the participation index for each pair of ROIs to belong to the same community.

The overlap between communities is evaluated using the binary matrices *M* obtained from each repetition of the Louvain algorithm, where the element *M_ij_* is 1 if the ROIs *i* and *j* belong to the same community and 0 otherwise. The overlap measure for two matrices *M* ^1^ and *M*^2^ is given by 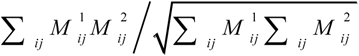, giving a number between 0 (no overlap) and 1 (identical matrices).

### Statistical Analyses

In order to assess to what extent a set of brain regions (such as the rich club) collectively exhibits significant differences from the cortex as a whole, we used a clustered-based bootstrapping procedure. That is, we averaged the measure of interest across regions falling within a previously defined cluster of size *N* (such as the rich club forming a theoretically motivated a-priori cluster). Subsequently, we repeatedly (10,000 times) sampled *N* regions from the entire set of cortical regions with replacement and calculated the average within each sample.

In order to assess differences observed for tasks with respect to rest, we employed a blocked bootstrapping procedure. Null distributions for each task were created from running connectivity optimization for 1,000 randomly resampled functional rest and task samples, calculating the difference (task - rest) for effective connectivity, and obtaining average changes in the measure of interest. Samples were created by first randomly drawing from the subject pool with replacement and then randomly placing one of each subject’s two states in the *rest* and the other in the *task* sample repeatedly until each sample comprised 14 subjects. Connectivity and local variability (sigma) were then optimized in each of these samples.

We compared the overlap of the communities corresponding the estimated ECs with that for random communities generated by partitioning the ROI indices into 4 to 6 groups (same range as the original communities). We repeated the process 1000 times to obtain a Null distribution of overlap measures. The difference between distributions was assessed using Welch’s t-test.

All p-values reported in the results have been corrected for multiple comparisons according to the false discovery rate (FDR) controlling procedure.

## Results

### Rich club

We followed the procedure outlined in Senden *et al.* (2017) to identify rich club regions in the SC. Specifically, we binarized the SC matrix by setting its non-zero entries to one. From this binary matrix rich club coefficients were calculated as the fraction of the number of existing connections between regions with degree larger than *k* to the possible number of connections among these regions (Colizza et al., 2006; Martijn P. van den Heuvel & Sporns, 2011; Zhou & Mondragon, 2004). Next, the statistical significance of rich club coefficients for each degree *k* was determined by calculating the rich club coefficients for a set of 1000 degree-preserving rewired adjacency matrices (Maslov & Sneppen, 2002) and identifying the first *k* for which the rich club coefficient of the binarized SC was larger than the 95^th^ percentile of the rich club coefficients corresponding to the rewired matrices. In our data, this was the case for *k* = 21. Seven candidate rich club regions were subsequently identified as those whose degree exceeded this cutoff. These included the bilateral precuneus, the bilateral superior frontal cortex, the bilateral superior parietal cortex, and the right insula. To ensure that these regions were not only individually rich but indeed also formed part of a dense club we evaluated the contribution of each region to the internal density of the set. To that end, we calculated internal density of the full set as well as of subsets resulting from leaving each candidate region out once. This procedure revealed that removal of the right insula lead to an increase in internal density by 21.33%. Since this exceeded the 95^th^ percentile of observed density changes (± 8.31%), the right insula was not considered a rich club region in this study. The six remaining regions have been identified as rich club hub regions in a multitude of studies (e.g. Daianu et al., 2015; Dennis et al., 2013; M. P. van den Heuvel, Scholtens, Feldman Barrett, Hilgetag, & de Reus, 2015; Martijn P. van den Heuvel & Sporns, 2011)

### Local variability

We first examined the local variability (Σ) exhibited by rich club and peripheral regions for resting and the four task states. Local variability exhibited by rich club regions was low compared to peripheral regions irrespective of state. Specifically, mean local variability within the rich club was 0.35 (95% CI [0.10, 0.59]), 0.40 (95% CI [0.19, 0.60]), 0.34 (95% CI [0.11, 0.57]), 0.39 (95% CI [0.24, 0.45]), and 0.38 (95% CI [0.16, 0.60]) for rest, the n-back task, the flanker task, the mental rotation task, and the odd-man out task, respectively. In contrast, local variability exhibited by peripheral regions was 1.36 (95% CI [1.14, 1.58]), 1.50 (95% CI [1.24, 1.77]), 1.43 (95% CI [1.19, 1.67]), 1.46 (95% CI [1.21, 1.70]), 1.41 (95% CI [1.16, 1.66]). A cluster-based bootstrapping test revealed that local variability exhibited by the rich club was significantly lower than that exhibited by peripheral regions in each state (figure 3A; p ≪ 0.01 for all comparisons). Note that these results do not imply that the model predicts no fluctuations in rich club activity as measured by the variance given by the diagonal of the model FC matrix 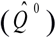. Mean variance for rich club regions was 1.00 (95% CI [0.78, 1.22]), 0.73 (95% CI [0.55, 0.91]), 0.80 (95% CI [0.63, 0.96]), 0.75 (95% CI [0.51, 0.99]), 0.77 (95% CI [0.61, 0.92]), for the five states, respectively. Mean variance of peripheral regions was 1.13 (95% CI [1.02, 1.25]), 0.93 (95% CI [0.83, 1.04]), 0.98 (95% CI [0.87, 1.09]), 0.91 (95% CI [0.80, 1.03]), and 0.93 (95% CI [0.82, 1.04]), for the five states respectively. Our clustered-based bootstrap procedures showed that differences between rich club and peripheral regions were not significant for any of the five states (figure 3B). Our model results thus reveal that fluctuating activity observed for the rich club is largely the result of network contributions from peripheral regions and not due to local variability.

**Figure 3:**
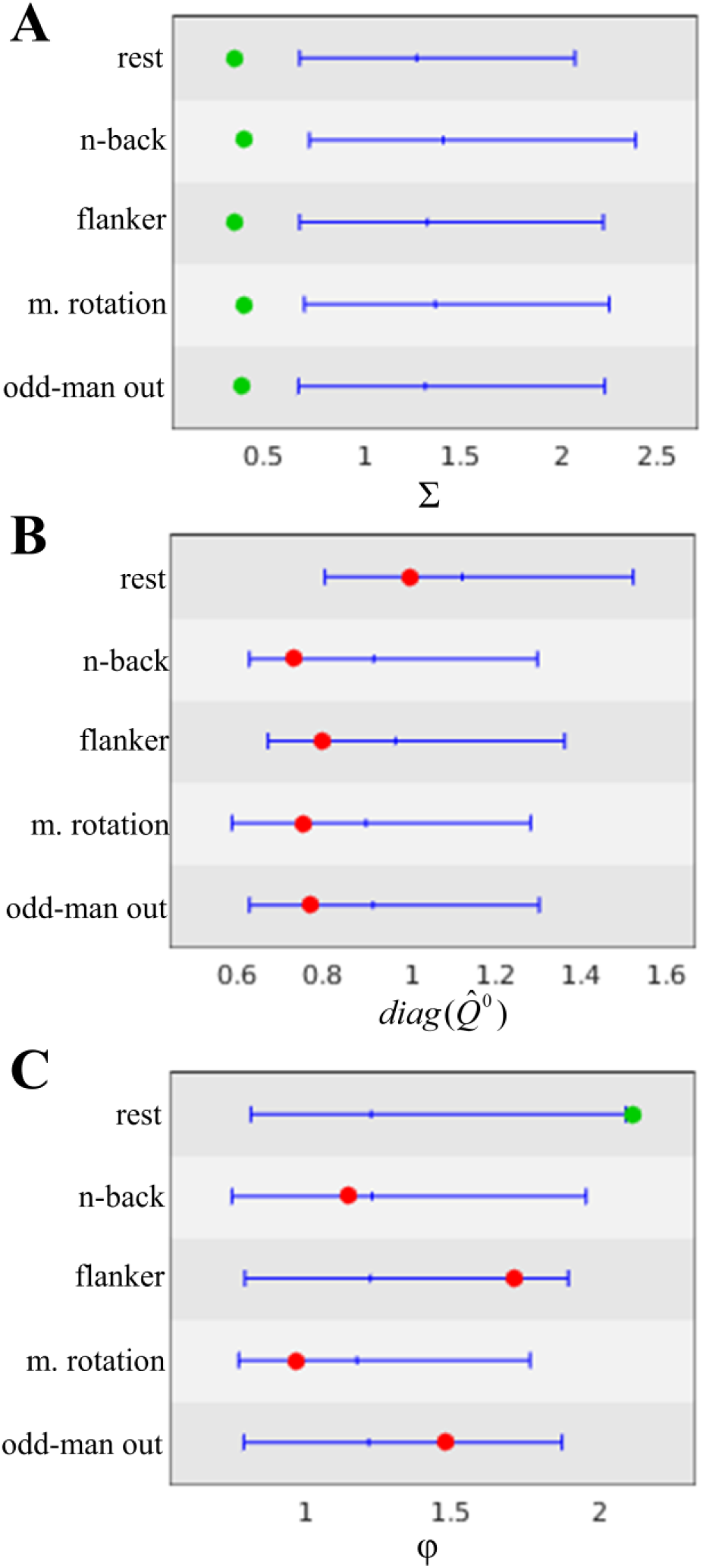
Cluster-based bootstrap analyses. This figure shows the region enclosed by the 2.5^th^ and 97.5^th^ percentile of sampled cluster averages (blue) for each task. Averages observed for the a priori rich club cluster falling inside this region (red) were considered non-significant while those falling outside this range (green) were considered significant. *Panel A)* shows results for local variability ∑. Local variability exhibited by the rich club was significantly lower than that exhibited by peripheral regions for rest and each task. *Panel B)* shows results for model variance *diag*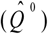. Variance observed for the rich club did not differ significantly from that observed for peripheral regions for either rest or task states. *Panel C)* shows results for the input-output ratio φ. The ratio exhibited by the rich club was significantly larger than that exhibited by peripheral regions for rest but not for any of the tasks

### Input-output ratio

We examined to what extent the rich club gates the input it receives from peripheral regions by computing the input-to-output ratio of total incoming/outgoing EC from and to the periphery. We focus on this ratio rather than absolute input and output strength since the rich club is likely to exceed peripheral regions in both simply by virtue of its high structural degree. The ratio provides an indication in how far the received input is passed on and is less dependent on structural degree. During rest, the total input the rich club receives from peripheral regions was roughly twice (φ_rc_ = 2.12, 95% CI [0.33, 3.90]) the output it sends to the periphery. In contrast, the ratio capturing in/out relationships among peripheral regions was only 1.14, 95% CI [1.03, 1.26]. During task performance, the rich club presented with lower input-output ratios. Specifically, the ratio exhibited by the rich club was 1.15 (95% CI [0.42, 1.88]), 1.71 (95% CI [0.52, 2.90]), 0.97 (95% CI [0.53, 1.41]), and 1.48 (95% CI [0.36, 2.60]) for the n-back, flanker, mental rotation, and odd-man out task respectively. The corresponding ratios observed for peripheral regions were 1.24 (95% CI [1.05, 1.44]), 1.18 (95% CI [1.03, 1.32]), 1.20 (95% CI [1.04, 1.36]), and 1.19 (95% CI [1.03, 1.34]). To assess whether input-output ratios exhibited by the rich club were significantly different from those exhibited by the cortex as a whole for each task, we again employed the cluster-based bootstrapping procedure. In line with our expectation, the input-output ratio was significantly larger for the rich club compared to peripheral regions during rest (p = 0.02) but not during any of the task states (figure 3C).

### Engaging in tasks is accompanied by increased rich club output

We follow up on the previous results by investigating to what extent decreases in the input-output ratio are due to the rich club receiving a decreased amount of input from the periphery or due to the rich club projecting an increased amount of output to the periphery during task states as compared to rest. To that end, we calculated the difference between EC observed during rest and during each of the task states and calculated the average change in input (output) received from (sent to) the periphery across the rich club. We assessed the significance of these changes per task by performing one-sided blocked bootstrap tests.

Input to the rich club from the periphery slightly increased (rather than decreased) for all tasks compared to rest. Specifically, average changes in input were 1.57 (95% CI [0.73, 2.41]), 1.45 (95% CI [0.67, 2.23]), 1.82 (95% CI [0.96, 2.67]), and 0.95 (95% CI [0.38, 1.52]) for the n-back, flanker, mental rotation, and odd-man out task, respectively. There could thus no significant decreases in input be observed. Increases in output from the rich club, on the other hand, were significant for all tasks except the flanker task (p = 0.04, p = 0.09, p = 0.04, and p = 0.04, for n-back, flanker, mental rotation, and odd-man out tasks). Specifically, change in output was 5.30 (95% CI [3.96, 6.91]), 5.71 (95% CI [4.72, 6.70]), and 3.73 (95% CI [1.97, 5.50]), for the n-back, mental rotation, and odd-man out task, respectively while it was 2.26 (95% CI [0.74, 3.79]) for the flanker task.

After establishing that the rich club is more conducive to whole-cortex communication during task performance compared to rest, we identified the specific peripheral targets of this increased output from the rich club. These are defined as those peripheral regions which receive an increase of output exceeding the 95^th^ percentile obtained from a blocked bootstrap procedure on differences in EC between tasks and rest from at least one rich club region. Figure 4 shows an overview of the thusly identified target regions per task. For the n-back task, the rich club displayed significantly increased output to 18 out of 62 peripheral regions. For the flanker task, it displayed significantly increased output to 11 peripheral regions. For the mental rotation task, it displayed significantly increased output to 18 peripheral regions. Finally, for the odd-man out task, it displayed significantly increased output to 17 peripheral regions. The increase in output strength was thus sparsely distributed. At the same time, the targets of rich club output for the different tasks were strongly overlapping. Specifically, the n-back task had 5 regions in common with the flanker task, 12 with the mental rotation task, and 10 with the odd-man out task. The flanker task had 9 and 8 regions in common with the mental rotation and odd-man out tasks, respectively. Finally, the mental rotation task had 13 regions in common with the odd-man out task.

**Figure 4:**
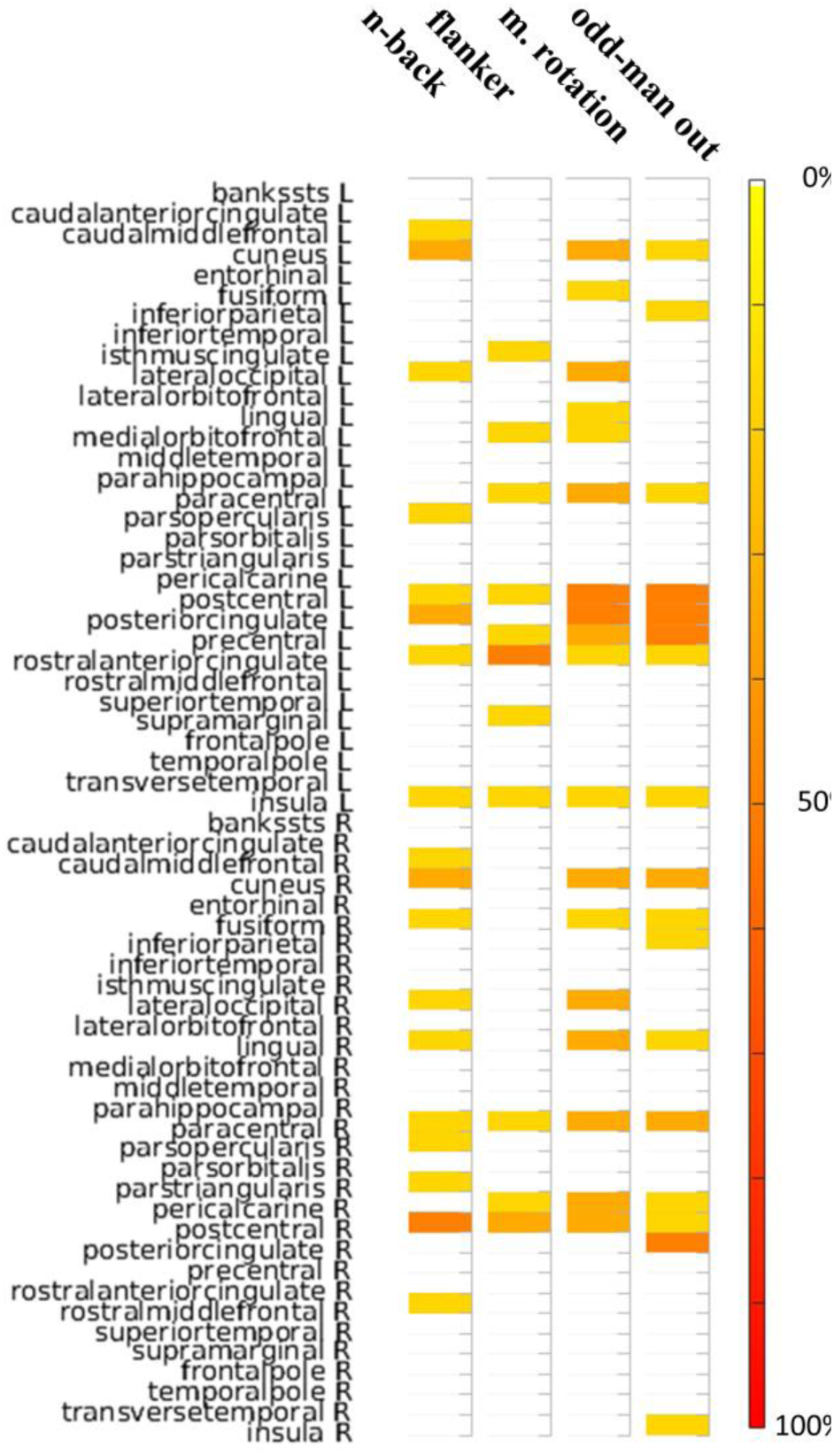
Targets of increased rich club output. This figure shows peripheral regions receiving increased output from at least one rich club region for each task as compared to rest. Colors indicate the percent of rich club regions sending increased output to a particular peripheral region with red reflecting a larger percentage. Targets of rich club output for the different tasks are largely overlapping. These regions include the bilateral cuneus cortex, the bilateral postcentral gyrus, the left rostral anterior cingulate cortex, the left insula, the right paracentral lobule, the left paracentral lobule, the left precentral gyrus, the left posterior-cingulate cortex, the right fusiform gyrus, the right lingual gyrus, and the right pericalcarine cortex. These regions are generally involved in processes common to each task such as visual perception, attention, and conflict monitoring (see discussion). At the same time, each task also presents with a set of regions serving as targets for increased rich club output uniquely for that task.

The pairwise analytical probability of obtaining the observed or more extreme degrees of overlap given the total number of regions selected per task constitutes a p-value for the Null hypothesis that the observed degree of overlap occurred by chance. FDR corrected p-values for each task pair are presented in table 2. As can be appreciated from this table, the degree of overlap was significant for all task pairs with the exception of the [n-back, flanker] pair. This indicates that many targets for the increased output of the rich club might be task-general. Indeed, the bilateral postcentral gyrus, the left rostral anterior cingulate cortex, the left insula, and the right paracentral lobule are targets of increased rich club output for all tasks. If regions which are target of increased output from the rich club for a majority of tasks are included, this set extends to include the bilateral cuneus cortex, the left paracentral lobule, the left precentral gyrus, the left posterior-cingulate cortex, the right fusiform gyrus, the right lingual gyrus, and the right pericalcarine cortex.

**Table 2:** Probability of observed overlap of target regions between tasks.

|  | n-back | flanker | mental rotation | odd-man out |
| --- | --- | --- | --- | --- |
| n-back |  | 0.17 | <0.01 | <0.01 |
| flanker | 0.17 |  | <0.01 | <0.01 |
| mental rotation | <0.01 | <0.01 |  | <0.01 |
| odd-man out | <0.01 | <0.01 | <0.01 |  |

In addition to these task-general target regions for rich club output, each task also presents with a specific set of target regions. For the n-back task these regions include the bilaretal caudal middle frontal gyri, the bilateral lateral occipital cortex, the bilateral pars opercularis, the right pars triangularis, and the right rostral middle frontal gyrus. For the flanker task target regions include the left isth^i^mus-cingulate cortex, the left medial orbitofrontal cortex, and the supramarginal gyrus. For the mental rotation task target regions include the bilateral lateral occipital cortex, the left fusiform gyrus, the left lingual gyrus, and the left medial orbitofrontal cortex. Finally, for the odd-man out task target regions include the bilateral inferior parietal cortex, the right insula, and the right posterior-cingulate cortex.

### Task-specific network communication

The final set of analyses focuses on task-related network communication as well as dynamics as compared to rest. In terms of communication, target regions of increased rich club output likely constitute the set of specialized peripheral regions whose integration into functional networks underlies task performance. As such, for each task we would expect higher levels of communication and hence larger empirical covariance between these target regions as compared to the remaining peripheral regions (i.e. those not forming targets of increased rich club output). Indeed, we find that average covariance (excluding variance) among target peripheral regions was 0.44 (95% CI [0.340, 0.47]), 0.42 (95% CI [0.35 0.47]), 0.48 (95% CI [0.43, 0.53]), and 0.48 (95% CI [0.44, 0.52]), for the n-back, flanker, mental rotation, and odd-man out task, respectively. For comparison, average covariance among nontarget peripheral regions was 0.30 (95% CI [0.29, 0.31]), 0.34 (95% CI [0.32 0.35]), 0.28 (95% CI [0.27, 0.29]), and 0.30 (95% CI [0.29, 0.31]), for the four tasks, respectively (see figure 6a). To assess whether average covariance among target peripheral regions was significantly higher than among non-target peripheral for each task, we again employed the cluster-based bootstrapping procedure with the set of target regions identified in each task as cluster and generating a Null distribution by sampling new clusters of equal size from all peripheral regions. In line with our expectation, average covariance was significantly larger for target compared to non-target peripheral regions for the n-back (p = 0.05), mental rotation (p = 0.04), and odd-man out (p = 0.04) but not the flanker task (p = 0.189).

At the same time, variance (and to a lesser extent covariance) decreased homogeneously across the cortex for task states compared to rest (see figures 5b and 6b). Specifically, average empirically observed variance across all cortical regions was 0.15 (95% CI [0.13, 0.16]), 0.12 (95% CI [0.11 0.13]), 0.13 (95% CI [0.11, 0.14]), 0.12 (95% CI [0.11, 0.13]), and 0.12 (95% CI [0.11, 0.13]), for rest and the four tasks, respectively. Differences in variance between task and rest were significant according to a blocked bootstrap test for the n-back (p = 0.04), mental rotation (p = 0.04), and odd-man out (p = 0.04) tasks while it marginally failed to reach significance for the flanker task (p = 0.053). This global reduction in variance can be explained by decreases in the measured time constants τ_x_ for individual regions in all tasks (see figure 5A; the ordering of regions for each task is given in table 3). These decreased nodal time constants lead to faster network dynamics (as measured by τ_c_, the negative inverse of the dominating eigenvalue of the Jacobian) and weaken recurrent network interactions. Non-significant increases in local variability ∑ and EC (figure 5B & C), in conjunction with significant, highly specific, increases in outgoing EC from the rich club (figure 6C), partially offset these decoupling effects and are likely necessary to sustain meaningful network communication in light of faster network dynamics during task performance.

**Figure 5:**
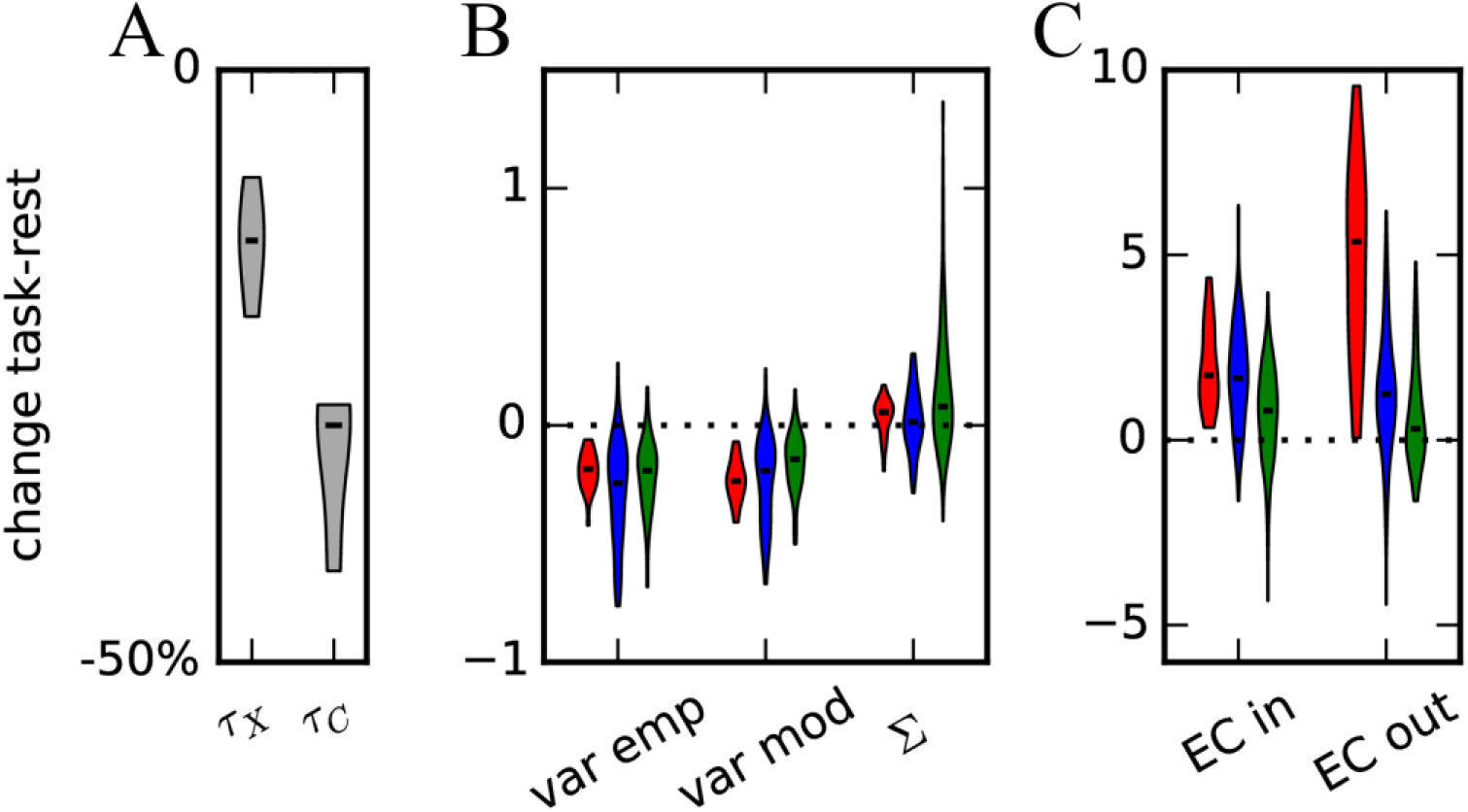
Changes in BOLD properties and model estimates with respect to rest. *Panel A)* shows changes in empirically measured region-specific and network time constants (τ_x_ and τ_c_, respectively) from rest to task averaged across tasks. Decreases for both constants reveal that cortical dynamics are generally faster during task performance as compared to rest. *Panel B)* shows changes in empirically observed variance, model variance, and local variability parameter Σ. Red indicate rich club regions, blue indicates peripheral regions serving as target for increased rich club in at least one task, and green indicates the remaining regions. *Panel C)* shows changes in both incoming and outgoing effective connectivity for the three groups of regions (color coding as before).

**Figure 6:**
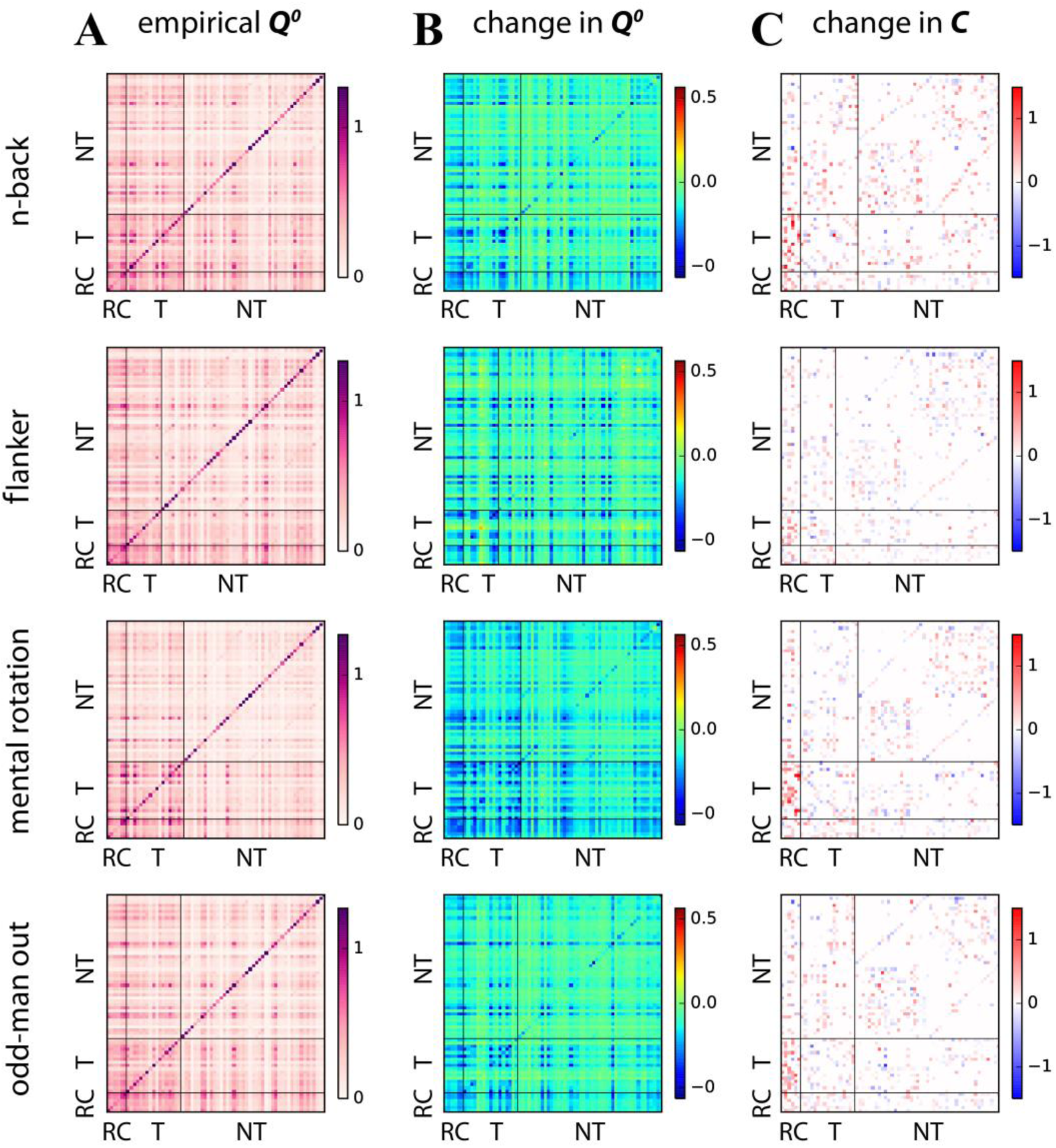
Comparison between whole-cortex connectivity observables and estimates for the four tasks. *Panel A)* displays empirical covariance matrices. The two horizontal and vertical lines demarcate rich club regions (RC) from peripheral regions serving as target in a specific task (T) and those, in turn, from the remaining peripheral regions (NT). See table 3 for a detailed overview of region order. *Panel B)* shows the change in covariance with respect to rest. *Panel C)* shows changes in effective connectivity with respect to rest.

**Table 3:**
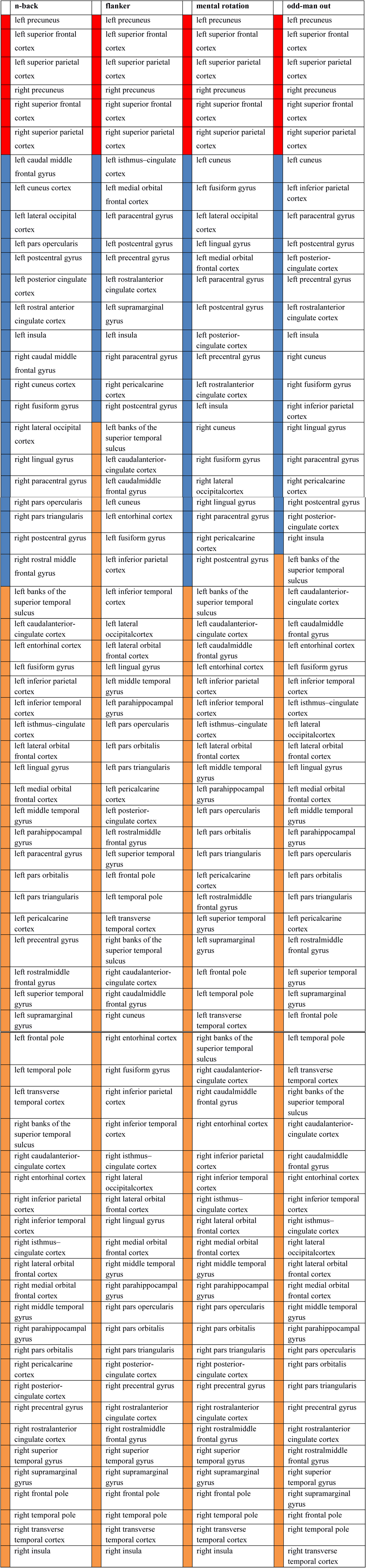
Order of regions in figures 6 and 7. For each task colors indicate whether a region belongs to the rich club (red), is a target peripheral region (blue), or a non-target peripheral region (green).

|  | **n-back** |  | **flanker** |  | **mental rotation** |  | **odd-man out** |
| --- | --- | --- | --- | --- | --- | --- | --- |
|  | left precuneus |  | left precuneus |  | left precuneus |  | left precuneus |
|  | left superior frontal cortex |  | left superior frontal cortex |  | left superior frontal cortex |  | left superior frontal cortex |
|  | left superior parietal cortex |  | left superior parietal cortex |  | left superior parietal cortex |  | left superior parietal cortex |
|  | right precuneus |  | right precuneus |  | right precuneus |  | right precuneus |
|  | right superior frontal cortex |  | right superior frontal cortex |  | right superior frontal cortex |  | right superior frontal cortex |
|  | right superior parietal cortex |  | right superior parietal cortex |  | right superior parietal cortex |  | right superior parietal cortex |
|  | left caudal middle frontal gyrus |  | left isthmus–cingulate cortex |  | left cuneus |  | left cuneus |
|  | left cuneus cortex |  | left medial orbital frontal cortex |  | left fusiform gyrus |  | left inferior parietal cortex |
|  | left lateral occipital cortex |  | left paracentral gyrus |  | left lateral occipital cortex |  | left paracentral gyrus |
|  | left pars opercularis |  | left postcentral gyrus |  | left lingual gyrus |  | left postcentral gyrus |
|  | left postcentral gyrus |  | left precentral gyrus |  | left medial orbital frontal cortex |  | left posterior-cingulate cortex |
|  | left posterior cingulate cortex |  | left rostral anterior cingulate cortex |  | left paracentral gyrus |  | left precentral gyrus |
|  | left rostral anterior cingulate cortex |  | left supramarginal gyrus |  | left postcentral gyrus |  | left rostral anterior cingulate cortex |
|  | left insula |  | left insula |  | left posterior-cingulate cortex |  | left insula |
|  | right caudal middle frontal gyrus |  | right paracentral gyrus |  | left precentral gyrus |  | right cuneus |
|  | right cuneus cortex |  | right pericalcarine cortex |  | left rostral anterior cingulate cortex |  | right fusiform gyrus |
|  | right fusiform gyrus |  | right postcentral gyrus |  | left insula |  | right inferior parietal cortex |
|  | right lateral occipital cortex |  | left banks of the superior temporal sulcus |  | right cuneus |  | right lingual gyrus |
|  | right lingual gyrus |  | left caudal anterior-cingulate cortex |  | right fusiform gyrus |  | right paracentral gyrus |
|  | right paracentral gyrus |  | left caudal middle frontal gyrus |  | right lateral occipital cortex |  | right pericalcarine cortex |

|  |  |  |  |
| --- | --- | --- | --- |
| right pars opercularis | left cuneus | right lingual gyrus | right postcentral gyrus |
| right pars triangularis | left entorhinal cortex | right paracentral gyrus | right posterior-cingulate cortex |
| right postcentral gyrus | left fusiform gyrus | right pericalcarine cortex | right insula |
| right rostral middle frontal gyrus | left inferior parietal cortex | right postcentral gyrus | left banks of the superior temporal sulcus |
| left banks of the superior temporal sulcus | left inferior temporal cortex | left banks of the superior temporal sulcus | left caudalanterior-cingulate cortex |
| left caudalanterior-cingulate cortex | left lateral occipitalcortex | left caudalanterior-cingulate cortex | left caudalmiddle frontal gyrus |
| left entorhinal cortex | left lateral orbital frontal cortex | left caudalmiddle frontal gyrus | left entorhinal cortex |
| left fusiform gyrus | left lingual gyrus | left entorhinal cortex | left fusiform gyrus |
| left inferior parietal cortex | left middle temporal gyrus | left inferior parietal cortex | left inferior temporal cortex |
| left inferior temporal cortex | left parahippocampal gyrus | left inferior temporal cortex | left isthmus–cingulate cortex |
| left isthmus–cingulate cortex | left pars opercularis | left isthmus–cingulate cortex | left lateral occipitalcortex |
| left lateral orbital frontal cortex | left pars orbitalis | left lateral orbital frontal cortex | left lateral orbital frontal cortex |
| left lingual gyrus | left pars triangularis | left middle temporal gyrus | left lingual gyrus |
| left medial orbital frontal cortex | left pericalcarine cortex | left parahippocampal gyrus | left medial orbital frontal cortex |
| left middle temporal gyrus | left posterior-cingulate cortex | left pars opercularis | left middle temporal gyrus |
| left parahippocampal gyrus | left rostralmiddle frontal gyrus | left pars orbitalis | left parahippocampal gyrus |
| left paracentral gyrus | left superior temporal gyrus | left pars triangularis | left pars opercularis |
| left pars orbitalis | left frontal pole | left pericalcarine cortex | left pars orbitalis |
| left pars triangularis | left temporal pole | left rostralmiddle frontal gyrus | left pars triangularis |
| left pericalcarine cortex | left transverse temporal cortex | left superior temporal gyrus | left pericalcarine cortex |
| left precentral gyrus | right banks of the superior temporal sulcus | left supramarginal gyrus | left rostralmiddle frontal gyrus |
| left rostralmiddle frontal gyrus | right caudalanterior-cingulate cortex | left frontal pole | left superior temporal gyrus |
| left superior temporal gyrus | right caudalmiddle frontal gyrus | left temporal pole | left supramarginal gyrus |
| left supramarginal gyrus | right cuneus | left transverse temporal cortex | left frontal pole |
| left frontal pole | right entorhinal cortex | right banks of the superior temporal sulcus | left temporal pole |
| left temporal pole | right fusiform gyrus | right caudalanterior-cingulate cortex | left transverse temporal cortex |
| left transverse temporal cortex | right inferior parietal cortex | right caudalmiddle frontal gyrus | right banks of the superior temporal sulcus |
| right banks of the superior temporal sulcus | right inferior temporal cortex | right entorhinal cortex | right caudalanterior-cingulate cortex |
| right caudalanterior-cingulate cortex | right isthmus–cingulate cortex | right inferior parietal cortex | right caudalmiddle frontal gyrus |
| right entorhinal cortex | right lateral occipitalcortex | right inferior temporal cortex | right entorhinal cortex |
| right inferior parietal cortex | right lateral orbital frontal cortex | right isthmus–cingulate cortex | right inferior temporal cortex |
| right inferior temporal cortex | right lingual gyrus | right lateral orbital frontal cortex | right isthmus–cingulate cortex |
| right isthmus–cingulate cortex | right medial orbital frontal cortex | right medial orbital frontal cortex | right lateral occipitalcortex |
| right lateral orbital frontal cortex | right middle temporal gyrus | right middle temporal gyrus | right lateral orbital frontal cortex |
| right medial orbital frontal cortex | right parahippocampal gyrus | right parahippocampal gyrus | right medial orbital frontal cortex |
| right middle temporal gyrus | right pars opercularis | right pars opercularis | right middle temporal gyrus |
| right parahippocampal gyrus | right pars orbitalis | right pars orbitalis | right parahippocampal gyrus |
| right pars orbitalis | right pars triangularis | right pars triangularis | right pars opercularis |
| right pericalcarine cortex | right posterior-cingulate cortex | right posterior-cingulate cortex | right pars orbitalis |
| right posterior-cingulate cortex | right precentral gyrus | right precentral gyrus | right pars triangularis |
| right precentral gyrus | right rostralanterior cingulate cortex | right rostralanterior cingulate cortex | right precentral gyrus |
| right rostralanterior cingulate cortex | right rostralmiddle frontal gyrus | right rostralmiddle frontal gyrus | right rostralanterior cingulate cortex |
| right superior temporal gyrus | right superior temporal gyrus | right superior temporal gyrus | right rostralmiddle frontal gyrus |
| right supramarginal gyrus | right supramarginal gyrus | right supramarginal gyrus | right superior temporal gyrus |
| right frontal pole | right frontal pole | right frontal pole | right supramarginal gyrus |
| right temporal pole | right temporal pole | right temporal pole | right frontal pole |
| right transverse temporal cortex | right transverse temporal cortex | right transverse temporal cortex | right temporal pole |
| right insula | right insula | right insula | right transverse temporal cortex |

Finally, we examined whether the changes observed in output connections across tasks have a homogeneous effect over the whole network. The goal was to test whether the network could be simply summarized into three groups (rich club, target and non-target regions) that clearly appeared in figure 6A-C. To do so, we performed community detection (see Methods) to partition the cortex into strongly connected subnetworks. This method compensates for the dense connectivity of rich club regions such that they do not belong to the same community. The outcome shown in figure 7A reveals reorganization of the communities for each task, both for target and non-target peripheral regions. To quantify the heterogeneities of the task-specific EC, we measured the overlap between partition matrices observed in the four tasks. Average partition matrices for the four tasks are displayed in figure 7A. Interestingly, the overlap between the task-communities displayed in figure 7B was at the same level as when randomly shuffling 20-50% of the ROIs from a given community (undistinguishable using Welch t-test with p>0.5). In comparison, shuffling only 10-20% of the ROIs results in much more overlap and shuffling all (random) in much less overlap, both with p ≪ 0.01. This indicates that changes in EC collectively induce a profound reconfiguration of communication at the network level, with an intermediate level of complexity.

**Figure 7:**
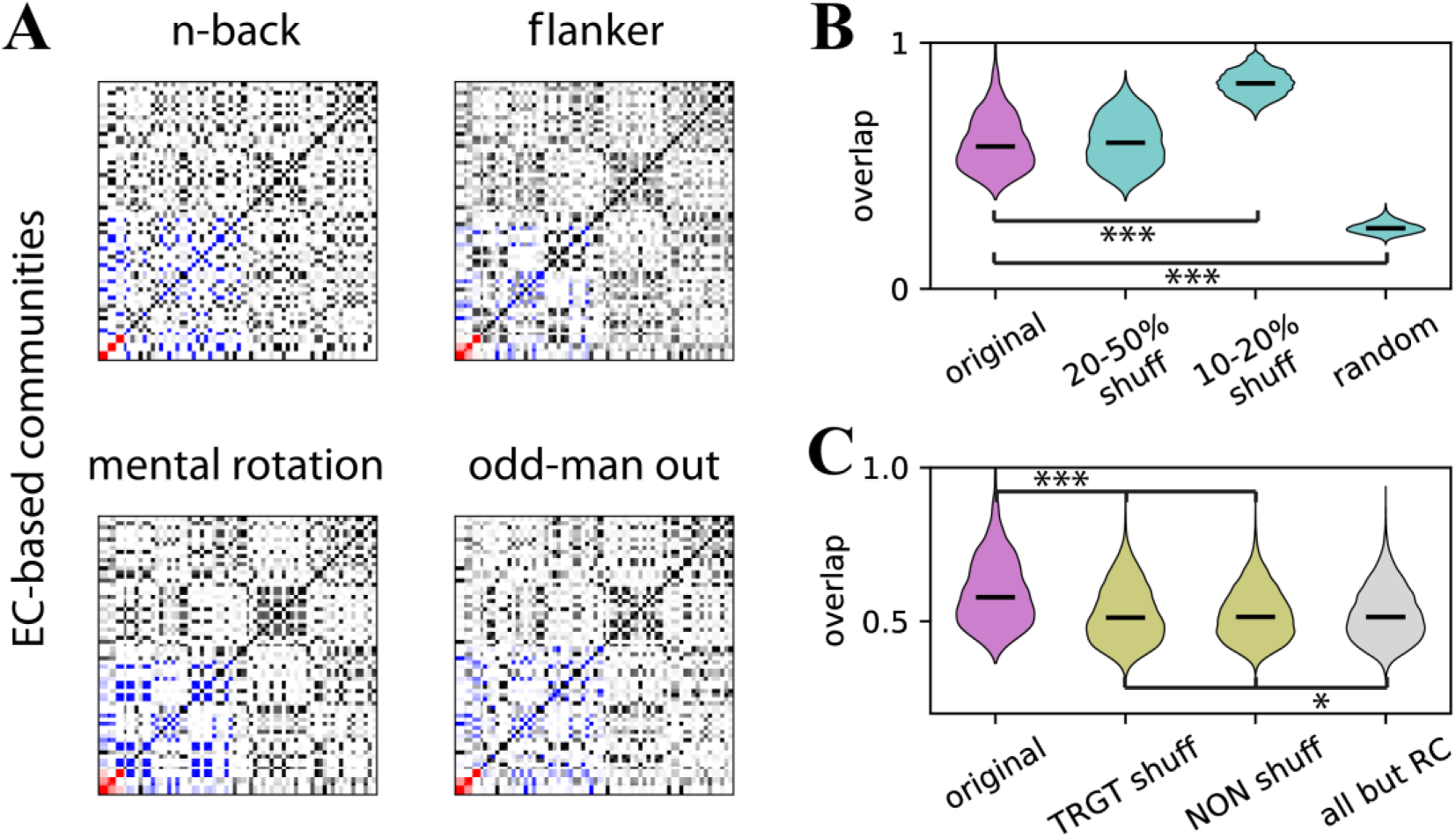
Overlap between task-related communities. *Panel A)* shows community participation indices between all pairs of cortical regions. Rich club regions are indicated in red (RC), peripheral regions serving as target in a specific task in blue (TRGT), and non-target peripheral regions in black (NON). The plotted participation indices correspond to averages over 30 repetitions of the stochastic detection algorithm per task; the order of the ROIs is the same for the four matrices. *Panel B)* compares the overlap between the communities for pairs of tasks (original) with the overlap between a given community partition and its shuffled version by permuting a portion of the ROIs (as indicated in the x-labels; random corresponds to 100%). *Panel C)* compares the overlap as in B with group-specific shuffling: within target and within non-targets (yellow distributions). In addition, the grey distribution corresponds to shuffling the same number of regions as for the yellow distribution on the left, but chosen at random from the set of 62 peripheral regions (i.e excluding the rich club).

Moving one step further, we examined the contribution of the two groups of peripheral regions (target and non-target) to the community structure across tasks. We shuffled ten peripheral regions either within a group (yellow violin plots) or irrespective of the groups (in gray). As shown in figure 7C, shuffling either within the group of target or non-target peripheral regions significantly reduced the overlap (with p ≪ 0.01). Importantly, the overlap observed when shuffling target regions does not differ from that observed when shuffling non-target regions (p = 0.2), though they each differ slightly but significantly from shuffling across groups (p = 0.023). The fact that shuffling target and shuffling non-target regions has an identical effect on overlap implies that, in contrast to the possibility that one group remains stable across tasks while the other exhibits extreme reconfigurations, both groups contribute equally to reconfigurations observed at the network level. This is relevant as it suggests that while largely the same peripheral regions serve as targets in all tasks, their communication yet differs between tasks.

## Discussion

In line with the hypothesis that the rich club might serve as a gated relay for whole-cortex communication, we found that during rest the rich club impedes transmission of activation between peripheral brain regions as total outgoing EC from the rich club was half of its total incoming EC. During task performance, on the other hand, a larger proportion of the input received by the rich club was passed on. The decrease in the input-output ratio observed for the rich club was not due to increases in local fluctuations as these remained consistently low irrespective of state. Neither was it due to decreases in input strength, the rich club always listens to the entire cortex. Instead, the change in input-output ratio was due to increased output strength from the rich club to the remainder of the cortex. Furthermore, outgoing effective connectivity did not increase for all regions, but only for a specific set of brain regions comprising between 18 and 29 percent of the periphery. These target peripheral regions showed strong coupling among each other (as measured by their empirical covariance) indicative of their functional relevance in their respective task. A subset of these peripheral regions served as targets of increased output from the rich club for all tasks while others were targeted specifically during individual tasks. The former subset included the cuneus cortex and the fusiform and lingual gyri which are involved in visual processing such as object recognition (e.g. Haxby, Gobbini, Furey, & Ishai, 2001; Ishai, Ungerleider, & Martin, 1999; Machielsen, Rombouts, & Barkhof, 2000; Mangun, Buonocore, & Girelli, 1998; Okada, Tanaka, Nakai, Nishizawa, & Inui, 2000; Vanni, Tanskanen, Seppa, Uutela, & Hari, 2001). It further included the posterior and anterior cingulate cortices which are involved in regulating the focus of attention and error/conflict monitoring, respectively (e.g. Bush, Luu, & Posner, 2000; Hampson, Driesen, Skudlarski, Gore, & Constable, 2006). Finally, it included the postcentral gyrus and the insula which are involved in sensing touch and motor control, respectively (e.g. Anderson, Jenkins, Brooks, & Hawken, 1994; Fink, Frackowiak, Pietrzyk, & Passingham, 1997). All of these functions pertain to each of our tasks as stimuli were presented visually, tasks were cognitively demanding, and motor responses were required.

Those regions serving as target for increased rich club output specifically for individual tasks have generally been associated with functions directly relevant for the task at hand. For the visual n-back task, regions included the caudal middle frontal gyri and the pars triangularis which have been implicated in visuospatial and semantic working memory as well as selection among competing items in memory (e.g. Carlson et al., 1998; Gabrieli, Poldrack, & Desmond, 1998; Leung, Gore, & Goldman-Rakic, 2002; Thompson-Schill, D’Esposito, & Kan, 1999). Other regions to which rich club output was increased during the n-back task were the lateral occipital cortex and the pars opercularis which have been associated with object recognition and selective response suppression (e.g. Forstmann, van den Wildenberg, & Ridderinkhof, 2008; Grill-Spector, Kourtzi, & Kanwisher, 2001). For the flanker task, target regions included the medial orbitofrontal cortex and the supramarginal gyrus; both related to decision making and evaluating decision outcomes (e.g. Kringelbach, 2005; Schoenbaum, Takahashi, Liu, & McDannald, 2011; Silani, Lamm, Ruff, & Singer, 2013). For the mental rotation task, regions included the lateral occipital cortex, the fusiform gyrus, and the lingual gyrus which are involved in encoding visual stimuli and object recognition (e.g. Grill-Spector et al., 2001; Haxby et al., 2001; Ishai et al., 1999; Machielsen et al., 2000; Okada et al., 2000) as well as the medial orbitofrontal cortex which is involved in decision making (e.g. Kringelbach, 2005). Finally, for the verbal odd-man out task, regions included the inferior parietal cortex implicated in word comprehension (e.g. Price, 1998) as well as the right posterior-cingulate cortex which is involved in regulating the focus of attention (e.g. Hampson et al., 2006).

In terms of whole-cortex network dynamics, we observed that BOLD variance decreased for task states as compared to rest and that these changes were due to faster cortical dynamics. This is in line with findings of He (2011) who showed that time-lagged autocorrelation of fMRI signals decreases during task performance suggestive of decreased long-range memory and more efficient online information processing. The fact that faster dynamics occur in spite of increases in recurrent EC further indicates that rest is closer to criticality than tasks. The cortex is thus optimally positioned in terms of its dynamics to allow for massive reorganization of functional interactions to meet task demands. This is also evident from our community detection analyses which revealed massive reconfigurations between tasks. Interestingly, reconfigurations occurred equally among target and non-target peripheral regions. This indicates that while many target peripheral regions appear to be relevant for all tasks and hence task-general in a univariate sense, their concrete interplay might nevertheless be highly task-specific. That is, different tasks (reflective of different cognitive domains) might not necessarily distinguish themselves in terms of which brain regions are involved in joint information processing but rather in terms of the communication channels established between these brain regions.

A number of points need to be considered when interpreting the findings of our study. First, we consider EC illuminating with respect to information flow. This rests on two assumptions; that our procedure is able to capture directional interactions between brain regions in the form of BOLD signal propagation, and that BOLD signal propagation reflects neural information flow. With regard to the former, our version of EC corresponds to the (linearized) interaction strengths that determine the BOLD activity propagation. Our results thus describe the net effects of changes in local activity and interactions in a phenomenological fashion. With regard to the latter, a recent study on mouse cortex has revealed that slow fluctuations in neuronal signaling (enveloping gamma-band activity) entrain vasomotion which, in turn, drives blood oxygenation (Mateo, Knutsen, Tsai, Shih, & Kleinfeld, 2017). Furthermore, it has been shown that the BOLD signal is closely related to perisynaptic activity in the form of the local field potential (LFP;Logothetis, Pauls, Augath, Trinath, & Oeltermann, 2001), at least as far as the neocortex is concerned (Ekstrom, 2009).

The LFP, in turn, reflects synaptic integration of information which can originate from local and remote sources (Herreras, 2016). Second, the noise diffusion model employed here provides a descriptive account of the BOLD signal rather than providing a mechanistic account of underlying neural signals. This is the appropriate level of detail since it captures those dynamics which can be observed with fMRI. Simulating the underlying neural signal would thus be more detailed but not necessarily more informative. In line with this, the noise diffusion model has previously been shown to recover the known effective connectivity in simulated neural signal stemming from a dynamic mean field model confirming that a more abstract description of the signal is sufficient (Gilson et al., 2016). For these reasons we are confident that we included all relevant details in our model while abstracting away from irrelevant ones (Boone & Piccinini, 2016). Third, we do not perform statistical analyses, and hence do not apply FDR-correction for multiple comparisons, when selecting target regions. Given that rich club output to the periphery increased significantly and that the system is closed, there must exist a set of target regions receiving this output. The 95^th^ percentile cutoff merely reflects a parameter used to identify these regions. Furthermore, analyses on target overlap across tasks shows that overlap is significantly larger than expected if regions had been selected at random within each task; i.e. if they constituted false positives. Fourth, we can currently not make any inferences regarding individual differences in whole-cortex EC. This would call for fMRI and diffusion-weighted MRI data to be acquired in the same subjects. Additionally, our sample size, while sufficient for analyses at the group level, is insufficient for characterizing this variability. Finally, it is known that parcellation schemes affect network parameters (Zalesky et al., 2010) rendering comparisons across studies difficult, if these use different schemes. Therefore, we relied on a widely used automated anatomical labeling template to increase comparability with previous studies. More research, using larger sample sizes and ideally limited to a single brain state, is thus necessary to investigate to what extent EC varies across subjects and how it depends on the choice of parcellation scheme.

Taken together, our results provide further support for the notion that the rich club serves as a central workspace wherein peripheral brain regions compete for the establishment of communication channels among them. During rest, slower, temporally dependent network dynamics reflect exploration of the brain’s functional repertoire. With impending task demands, network dynamics become faster in order to process information effectively. At the same time, the rich club increases its outgoing effective connectivity to establish communication among task-relevant peripheral regions and prevent global decoupling (a consequence of faster dynamics). With a transition to task performance, the rich club thus supports the interplay of a set of task-relevant peripheral regions as required for higher cognition.

## Acknowledgements

Authors MS and RG were supported by the European Research Council under the European Union′s Seventh Framework Programme (ERC-2010-AdG, ERC grant agreement no. 269853) and by the European Union’s Horizon 2020 research and innovation programme under grant agreement n. 720270 (HBP SGA1). Author GD was supported by the ERC Advanced Grant: DYSTRUCTURE (n. 295129), by the Spanish Research Project PSI2016-75688-P (AEI/FEDER), and by the European Union’s Horizon 2020 research and innovation programme under grant agreement n. 720270 (HBP SGA1). Author MG was supported by a Marie Sklodowska-Curie Action grant (H2020-MSCA-656547), by European Research Council Advanced Grant DYSTRUCTURE (Grant 295129), and by the European Union via the Human Brain Project (grant FP7-FET-ICT-604102). Author MPvdH was supported by a VIDI grant of The Netherlands Organization for Scientific Research (NWO 452-16-015), by a MQ Fellowship and by NWO ALW open (ALWOP.179). The authors declare no competing financial interests.

